# Buoyant-Antigen-Magnetic (BAM) ImmunoSeparation, isolation, and detection of Specific Pathogenic Bacterial Cells

**DOI:** 10.1101/2022.11.27.517249

**Authors:** Brandon H. McNaughton, Jeffrey N. Anker, Päivö K. Kinnunen

## Abstract

Separating, collecting, and detecting specific cells and molecules from complex and heterogeneous samples is key for many biomedical, environmental, and food safety applications. Antibodies are widely used to specifically bind cells and molecules to a solid support such as a column or suspended bead to separate, wash, and concentrate them. However, nonspecific binding of non-target cells and molecules to the support limits the specificity and resulting purity of traditional separation methods. Herein we describe a buoyant-antigen-magnetic (BAM) separation approach to increase specificity using two distinct separation processes with “AND” logic. Buoyant silica microbubbles and magnetic microbeads, each with their own antibody, bind to the target cell forming a BAM sandwich complex which can be separated from non-targeted cells (which are neither magnetic nor buoyant), as well as from cells bound to only single buoyant or magnetic microparticles but not both. *Escherichia coli* O157:H7, a pathogenic bacteria that can contaminate food products, was used as a model organism to establish a nearly 100-fold reduction in the non-specific carryover of non-target *Escherichia coli* bacteria in comparison to simple magnetic separation. Beads with no attached analyte were also removed. Furthermore, the BAM complexes could be visualized by eye (or camera) providing a rapid and sensitive method for pathogenic bacterial detection. Bacterial concentrations as low as 5 × 10^4^ CFU/ml were evident by eye and could easily be distinguished from non-target *S. aureus* bacteria even at much higher concentrations (5x10^7^ cfu/mL). This approach enabled simple user-friendly measurement for on-site detection and isolation without expensive equipment.

**HIGHLIGHTS:** 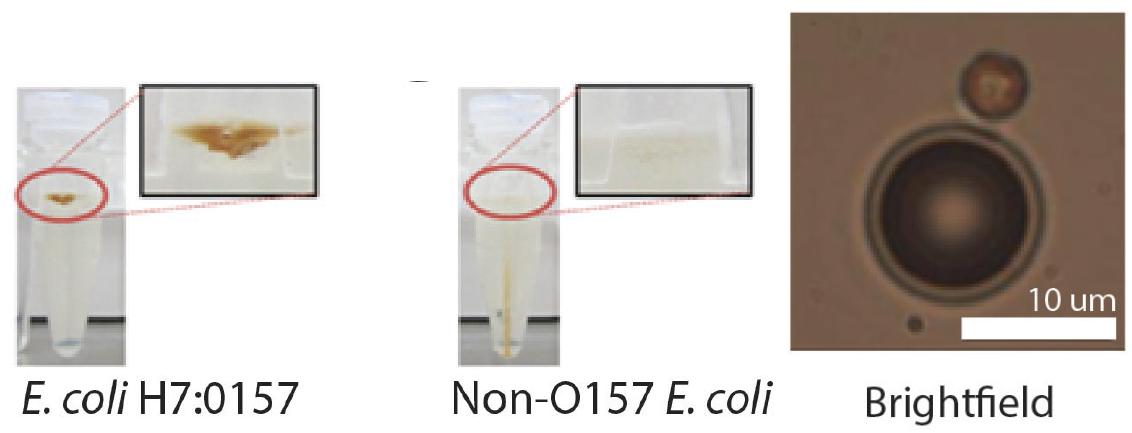

## 1.0 Introduction

In most applications, detecting and quantifying specific cells and molecules is a two-step process involving a sample preparation followed analysis. Improvements in analyte detection may come from enhancements to either sample preparation or detection. Instrumentation improvements, while sometimes dramatic, often face real-world performance limits due to inadequacies in sample preparation— for example, sophisticated analysis methods often require samples to be challengingly pure prior to measurement and less sophisticated instruments may have limited sensitivity. Thus, simple sample preparation steps that increase the sample purity and concentration prior to analysis can significantly improve detection.

An important approach to analyte cleanup has been the use of magnetic nanoparticles or microparticles with an affinity for the target molecule or cell. This strategy allows both concentration of target analytes from complex samples and removal of interfering substances that may affect downstream detection and measurement instruments. Indeed, analyte separation with magnetic beads is a key element in applications as varied as high throughput gene sequencing [1], molecular diagnostics in medical microbiology [2] and virology [3], cell-based immunologic and cancer tests [4], and food pathogen testing [5]. Since this technique was introduced over thirty years ago, commercially available magnetic bead methods have become commonplace in industrial and medical life science laboratories that isolate cells or macromolecules from complex samples [3], [4], [6], [7], [8].

While magnetic separation has improved research and clinical diagnostics, the method is not without limitations. A key shortcoming of magnetic separation is that the specificity of the method is governed by the performance of the bead functionalization. For cell capture and isolation, magnetic beads typically are coated with an antibody directed at a single target epitope which can lead to significant non-specific binding, especially when background cell concentrations are high. This problem arises in a manner analogous to the low specificity encountered in enzyme immunoassays based on analyte detection with a single antibody. Another limitation arises because the number of magnetic beads is conserved in the concentration process. The number of magnetic particles used in a separation process can be three or four orders of magnitude higher than the number of target cells—for example, in a food pathogen testing protocol one might use 10^7^ magnetic beads in a 1 mL volume that has 10^4^ cfu/mL concentration of bacteria. In practical terms that means for every magnetic bead that engages a target cell, there are 999 beads that are ‘empty’. Using standard techniques, these empty magnetic beads cannot be separated from beads carrying the desired payload—and unavoidably carry forward unwanted cells or inhibitors of detection methods, such as polymerase chain reaction. Ultimately, the presence of these empty beads and their non-target cargo affect downstream methods through lost specificity or poisoning of analytical chemistry. In applications like food testing, where samples can be highly complex (e.g., raw ground meat), non-specific binding is a major concern that limits the utility of current bead-based methods.

To date, improvements in magnetic-bead-based sample capture—including microfluidic approaches [9]–[11], formation of magnetic holes [12], particle sieving [13], or the substitution of magnetic particles with buoyant particles [14]–[22]—have not overcome the problems of low specificity and non-target carryover. Thus, in technical fields where magnetic bead technology is employed, there remains an unmet need for a sample preparation process with improved specificity. For molecular detection applications, double labelling molecules of interest can provide very high sensitivity and specificity, for example, Walt and co-workers combined microbead separation with fluorogenic enzymes for single molecule counting[23], [24], and Thraxton and co-workers combined magnetic particle labeling with DNA-functionalized gold nanoparticles for atomolar detection of prostate specific antigen[25], however, the readout mechanisms are relatively complex (involving microscopy and fluorescence) and the second label serves only for observation not of separation and downstream processing.

In this report, we describe the use of a pair of bead types—one that is magnetic and one that is buoyant—to achieve high specificity isolation of bacteria. The separation is based target cells binding to both an antibody-labeled magnetic bead and an antibody-labelled buoyant bead to form a suspended buoyant-antigen-magnetic (BAM) complex. By employing both magnetic and buoyant forces, only beads linked via the target cell having both buoyant and magnetic properties are carried through the separation process, as shown in Figure 1. We focus our attention on the recovery of specific capture of the human pathogen *E. coli* O157:H7 in comparison to non-specific capture of bacteria.

**Figure 1.**
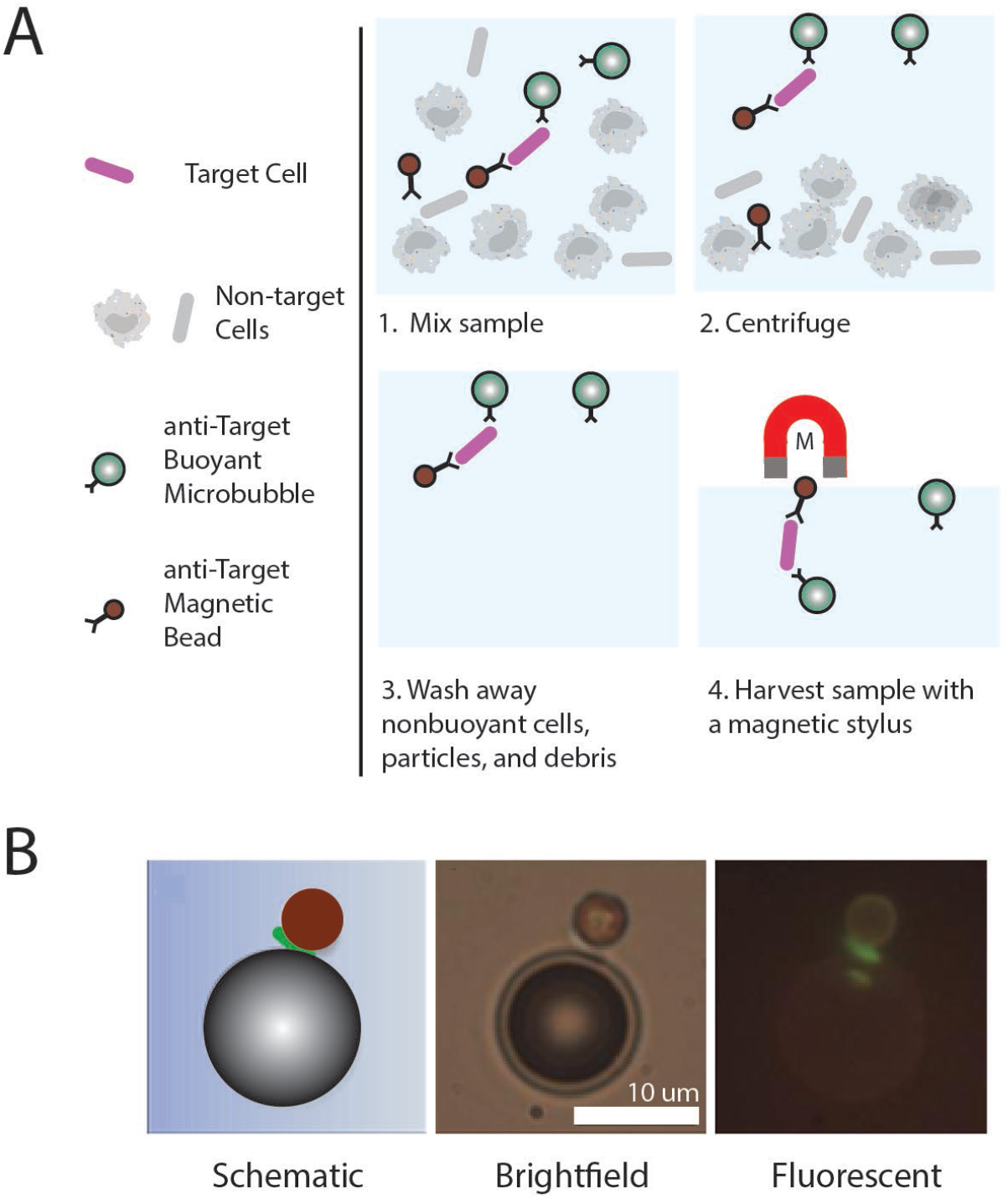
Operating principle of target cell capture using two-bead separation. Panel A. Typical workflow. The strategy relies on the formation of a suspended complex of a target cell with buoyant and magnetic particles. This permits the capture of particles meeting both magnetic and buoyant requirements in a manner analogous to sandwich immunoassay methods. Panel B: Example two-bead complex incorporating a silica microbubble and magnetic bead each surface functionalized with an antibody to *E. coli* H7:0157 and an interposed bacterium.

## 2.0 MATERIALS AND METHODS

### 2.1 Bacterial strains and methods

*E. coli* O157:H7 (ATCC 35150), non-O157 *E. coli* (ATCC 25922), and *Staphylococcus aureus* (ATCC 29213) were obtained from ATCC Microbiology (www.atcc.org). Cells were maintained cryopreserved until use. Mueller-Hinton agar and cation-adjusted Mueller-Hinton broth (CA-MHB) were used as growth media in all cases. Quantitative culture was performed using serial dilution and colony counting on agar plates, and all methods were in compliance with Clinical and Laboratory Standards Institute microbiological methods (clsi.org).

### 2.2. Magnetic bead capture

In experiments examining only magnetic bead performance, bacteria were captured using commercially available beads with their associated magnetic separator device (anti-*E. coli* O157 Dynabeads^®^, DynaMag™ -2 magnet, Life Technologies) following the manufacturer’s guidelines for use. The exact specification of the Dynabeads are proprietary, but previous studies have shown they are 2.8 μm in diameter and have a density of 1.3 g/cm^3^ In brief, 1 ml of bacterial suspension was mixed with 20 μl of functionalized magnetic beads (∼2x10^5^ beads) for 10 minutes at 37 °C. Samples then underwent three rounds of washing prior to bead collection and quantitative culture.

### 2.3 BAM separation

Silica glass microbubbles (3M) were functionalized with anti-*E. coli* O157:H7 antibodies (Bactrace®, KPL, www.kpl.com) using isoelectric point surface adsorption, washed, then resuspended to a final concentration of 10^7^ beads/ml. To bacterial suspensions were added 20 μl anti-*E. coli* O157 magnetic beads (∼2x10^5^ beads) and 100 μl anti-*E. coli* O157 buoyant beads (∼10^6^ beads). These were placed on a rotating end-over-end mixing platform at 60 rpm and 37°C for 15 minutes. The samples were then centrifuged for 2 minutes at 400 × g. The jointly buoyant/magnetic beads were then extracted with a magnetic stylus (PickPen, Cat # 23001, Sunrise Science Products, Inc.) and released into growth media for quantitative culture. See Figure 1 Panel A for an overview of the process.

### 2.4 Modified BAM separation for direct visualization of complexes

In a related set of experiments, the order of complex isolation was reversed such that the magnetic field was applied and released before complexes were allowed to float. See Figure 3 Panel A for an overview of the process.

### 2.5 Quantification of magnetic bead and BAM separation processes

Method performance was measured in two ways. First, the fraction of bacteria recovered, *f*, for each strain by each method was determined by quantitative culture, where:

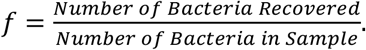

Typically, the total bacterial count in a sample was on the order of 5 × 10^7^ cfu in one milliliter of liquid. The relative affinity of either the magnetic or the two bead process for *E. coli* O157:H7 versus one of the two other test strains (non-O157 *E. coli* and *S. aureus*), was defined as:

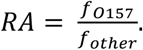

Differences in recovered fraction or relative affinity were compared using two sided t-tests on log-transformed data. All analysis was conducted in R 3.0.2.

## 3.0 RESULTS

### 3.1 Use of BAM separation for specific pathogenic bacteria recovery

As described in Section 2.3, the simultaneous mixing of bacteria and antibody-functionalized magnetic and buoyant silica microparticles led to the capture of *E. coli* O157:H7 bacteria into complexes that were jointly magnetic and buoyant (Figure 1). Quantitation of this phenomenon revealed that in general, magnetic-only separation resulted in larger capture efficiency than two-bead separation (Figure 2 Panels A and B), but with a significantly poorer specificity. The relative affinity for two bead complexes (Figure 2 Panel D) showed that capture of target *E. coli* and rejection of non-target *E. coli* or *S. aureus* was dramatically superior. In comparison to *S. aureus*, the relative affinity of the two-bead separation process for *E. coli* O157:H7 was almost 1,000-fold superior to magnetic-alone separation.

**Figure 2.**
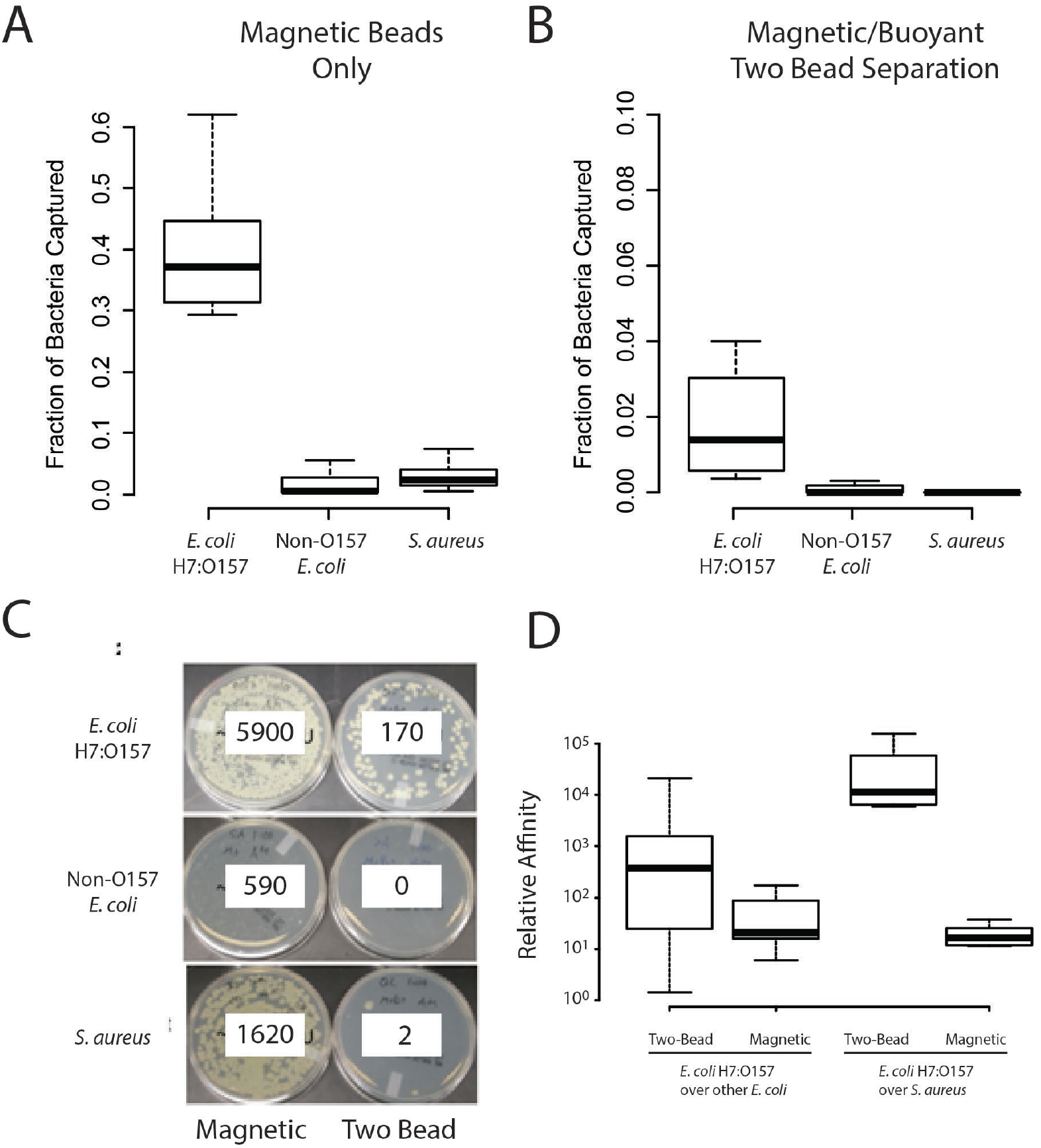
Performance of two-bead separation in comparison to standard magnetic bead method. Panels A and B: Fractional recovery of bacteria from cultures of *E. coli* O157:H7, a non-O157 *E. coli*, and *S. aureus*. In each case, beads were mixed with ∼10^7^ cfu of bacteria in a one milliliter volume. The fractional recovery represents the portion of all exposed bacteria recovered by each method. Recovery of non-O157 *E. coli* or *S. aureus* represents non-specific binding. The two-bead method recovered a significantly smaller fraction of available bacteria (p < 0.05 by t-test). Panel C. An example of raw culture data from these experiments. Panel D. The relative affinity of the magnetic and two-bead methods for *E. coli* O157:H7 (see text for details). Although the two-bead method isolated fewer target organisms in total (as in panels A and B), its specificity for the O157:H7 strain was remarkable – over ten-fold against non-O157 *E. coli* and over 1,000-fold against *S. aureus*. Data shown as 5^th^, 25^th^, 50^th^, 75^th^, and 95^th^ percentile box plots, n = 7 per condition.

### 3.2 Direct visualization of BAM complexes

By reversing the order of separation such that the magnetic field was applied before the buoyant force, BAM complexes could be directly visualized floating to the top of the test tube (Figure 3). This material appeared as a floating brown band at the top of the sample (due to the brown color of the magnetic particles), which was easily seen with *E. coli* O157:H7 but was not visible in samples containing either non-O157 *E. coli* or *S. aureus*. This is distinct from what occurs when the target is not present—and what happens in typical magnetic separation—where the magnetic beads sink to the bottom of the vial. Subsequent testing revealed that the floating brown band was visible down to a target analyte concentration of 5 × 10^4^ cfu/ml (Figure 3 Panel B).

**Figure 3.**
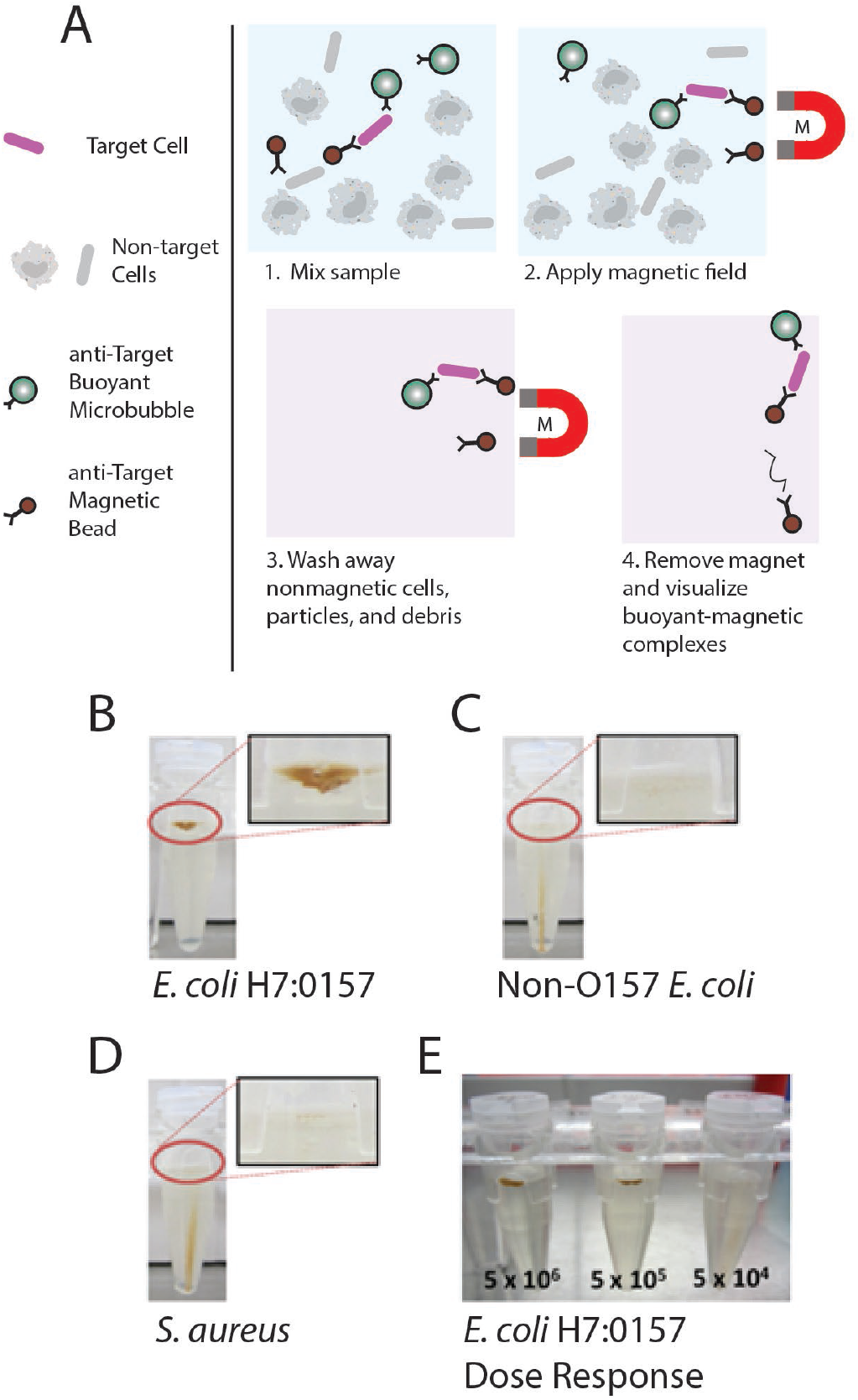
Alternative workflow that allows direct visualization of two-bead complexes. Panel A. Modified workflow from Figure 1 that performs magnetic separation first, followed by buoyant separation. Panels B, C, and D: Results of two-bead isolation showing buoyant brown material in the presence of *E. coli* O157:H7 but not in the presence of non-O157 *E. coli* or *S. aureus*. Given the high relative density of the brown magnetic particles relative to the aqueous test fluid, flotation of magnetic beads indicates simultaneous binding of buoyant microbubbles. Panel E: Level of sensitivity of this method for *E. coli* O157:H7, indicating that the floating two-bead complexes are visible to the unaided eye at a concentration of 5x10^5^ particles.

## 4.0 Discussion

Here we report the joint application of buoyant and magnetic microparticles to capture with high specificity pathogenic *E. coli O157:H7* from a liquid sample. By functionalizing the surfaces of glass microbubbles and magnetic particles with O157-specific antibodies, we created a fluid-suspended equivalent of a sandwich immunoassay. Compared to bacterial recovery with magnetic beads alone, we were able to achieve between 50- and 1,000-fold improved specificity over magnetic beads, as measured by quantitative culture. This improvement required neither a longer incubation time nor the introduction of new equipment.

We believe this approach may offer significant benefits in applications seeking to isolate bacteria, other cell types, or subcellular structures from complex backgrounds. Current methods in this area include fluorescence activated cell sorting (FACS) or a number of variations on bead-based capture (typically magnetic activated cell sorting, or MACS). Both technologies have important benefits and limitations. FACS is by far the most quantitative, with the separation process coupled to detailed characterization of surface and intracellular features of every particle sorted. Its limitations include its substantial costs for entry and maintenance as well as its slow speed and incapacity for high throughput sample handling. Bead-based MACS overcomes these limitations. However, magnetic based approaches suffer from low specificity (due to the use of one antibody type) and challenges in sample scale up due to the relationship between magnetic field strength and distance.

A two-bead approach may provide an economical middle ground —much greater target specificity can be achieved, and the application of magnetic field can be optimized by first spatially concentrating sample targets by using their buoyancy. As we show in Figure 3, the combination of buoyancy with the natural brown color of magnetic beads allowed for a simple and direct visualization of the target capture. Further modifications of the beads (e.g., fluorescence labeling of one of the components) may allow for highly sensitive visual detection of small numbers of target cells. This raises the possibility of application in point-of-testing technologies in the life sciences, industry (e.g., food production), and medicine.

The approach is appropriate for detection in resource-limited environments which are described in the World Health Organization’s ASSURED criteria (affordable, sensitive, specific, user-friendly, rapid and robust, equipment-free and deliverable to end users). The main equipment it needs is a magnet and potentially a camera, with disposables being buoyant microbubbles and magnetic microbeads, vials, and pipettes. The simplicity of the device and protocol would make it ideal for on-site analysis.

The speed of the separation depends upon the incubation time and particle concentration, as well as the buoyancy and magnetophoretic forces and distance the microbeads need to travel. Under the conditions we used, the magnetic separation occurred in <1 min and the following buoyant separation also occurred in <10 min when left to settle under gravity. In general, the external magnet is sufficient to saturate the magnetism of the superparamagnetic microbeads, and the separation then depends upon the magnetic field gradient (which is around ∼100 T/m near the permanent magnet) and particle diameter and is given by: v = d^2^ m_s_*∇H/18 η, where v is the velocity in m/s, d is the particle diameter (3x10^−6^) m, m_s_ is the saturation magnetization (theoretically around 29 kg/m for a particle with 1.3 g/cm^3^ comprised of 6% volume magnetite particles, 480 kg/m) and and 94% polystyrene), and η is the fluid viscosity (1.0x10^−3^ Pa-s for room temperature water) [30]. The buoyant rising velocity is given by (ρ_bead_− ρ_solution_)*g* d^2^ /18 η, where the buoyant bead density, ρ_bead_, is 600 kg/m^3^, the solution density, ρ _solution_, is ∼ 1000 kg/m^3^, g=9.8 N/kg under normal conditions, and d ranges from 10 μm to 30 μm. Overall, when the magnet is present, the magnetophoretic forces from a single 3 μm magnetic bead in a BAM complex were more than sufficient overcome the buoyancy force when the magnet is present, and when the magnet is released, single BAM particles will rise at ∼1-10 mm/min. This rising rate increase when many microbubbles are close to each other, as they pull surrounding fluid with them as they rise reducing the overall resistance and causing the ensemble to rise faster overall. Speed can also be increased via centrifugation. However, for many applications, waiting 10 minutes for low concentrations of buoyant beads to rise ∼ 1 cm under gravity would be acceptable.

An important limitation of the two-bead method is that we observed lower total cell recovery achieved. As shown in Figure 2 panels A and B, magnetic beads alone captured 38% of bacteria in the test sample, while buoyant-antigen-magnetic sandwich complexes captured 15% of the total bacteria. The severity of this limitation varies by application.

Where highly reliable detection is most important, decreased total yield may be acceptable given the much greater rejection of non-target analytes. In other applications, such as harvesting stem cells for therapeutic use, total yield may be of greater concern. We also expect that the performance of the BAM method can be improved by optimizing the particle concentration, particle size, sample volume, choice of complementary antibodies, order or number of steps (e.g., use of a buoyant microbubble separation followed by addition of concentrated magnetic particles for efficiently form BAM complex formation), and incubation time. Indeed, recent work with commercialized Akadeum microbubbles show >99% capture efficiency for red blood cells and circulating tumor cells,[26], [27].

## 5.0 Conclusion

In conclusion, we demonstrated highly specific capture, detection, and isolation of pathogenic bacteria using the BAM approach. This work addresses a pressing need for improved sample preparation in a variety of fields, by reducing non-specific binding, allowing immediate direct visualization of the separation process, and potentially expanding the volume of samples which can be processed with bead-based methods. Future work will study the reaction conditions to optimize separation sensitivity and specificity. We will also extend the approach to other pathogens, cells (e.g., circulating tumor cells), and analytes. To increase assay sensitivity and study specific and non-specific binding, we will image BAM complex as they rise due to buoyancy or are pulled to a magnet and optimize surface chemistry and incubation buffers.

## Acknowledgements

This work was supported in part by grant DMS 1224844 from the National Science Foundation (jgy). The authors would like to thank those that supported the early and ongoing development of this technology, including Sundu Brahmasandra, Elizabeth Craig, Maureen Carey. We would also like to thank the Coulter Foundation and MIIE for supporting the ongoing efforts of developing this work. The authors also thank Prannda Sharma for diligently working on the ongoing research efforts and Prof. Jouko Niinimaki. For additional information about two-bead separation, including videos and additional content, please see www.akadeum.com.

